# Tumour metabolic heterogeneity discrimination by video thermometry and concanamycin-sensitive proton pump measurements: Old tools integrated into advanced preclinical studies

**DOI:** 10.64898/2026.03.06.708889

**Authors:** Paula Gebe Abreu Cabral, Sávio Bastos de Souza, Brunna Xavier Martins, Scarlath Ohana P. dos Santos, Silvia M. R. Cadena, Tainara Micaele Bezerra, Raul F. Arruda, Renato M. da Silva, Sheila P. F. Cabral, Hassan Jerdy, Gustavo D. A. Braga, Luciana M. Mell, Raphael W. F. Menassa, André Lacerda de Abreu Oliveira, Arnoldo Rocha Façanha

**Author notes:** These authors contributed equally to this work (Paula Gebe Abreu Cabral, Sávio Bastos de Souza, Brunna Xavier Martins).

## Abstract

Phenotypic, genetic and metabolic tumour heterogeneity is a major factor contributing to cancer progression, metastasis, and resistance to therapy. Distinctive thermal and proton flux patterns can occur in cancer cells as a function of variations in the metabolic rate of the tumour mass, tumour margins, and normal tissues, which can be detected by video thermometry (VTM) as well as by Scanning Ion-selective Electrode Techniques (SIET). This study presents distinct thermal patterns associated with tumour heterogeneity observed in canine mammary cancer using VTM and investigate the correlation between these thermometric signatures with metabolic changes related to V-ATPase activity by comparing real-time VTM data with that from concanamycin-sensitive ATP hydrolysis and cell proton flux measurements. The results demonstrate that integrating SIET and VTM data sets can reveal metabolic signatures to assist the diagnosis, surgery and therapeutic monitoring. Considering the breast cancer hallmarks conservation in human, canines and other mammals, this study provides a first bioenergetic proof-of-concept for the potential of the integration of the VTM technology with enzymatic and electrophysiological analyses sensitive to concanamycin for the development of more effective diagnostic, prognostic and therapeutic approaches in veterinary as well as in preclinical medical studies.

## INTRODUCTION

Breast cancer is a prevalent malignancy characterized by uncontrolled growth of abnormal cells in the mammary tissue, forming tumours potentially fatal if not early detected and properly managed. Various types of tumours can arise within the mammary glands of most mammals, with a high incidence in both women and female dogs, with specific aggressiveness largely determined by their metabolic activity that supports the dissemination of metastatic cells, a primary cause of mortality (Bergers & Fendt, 2021). Tumour heterogeneity also includes genetic, phenotypic, and functional variations among cancer cells within a single tumour, as well as between distinct tumours in the same patient or among different patients (Turashvili & Brogi, 2017).

Diagnosing breast cancer accurately and promptly is still a challenge task. Even with improvements in mammography, the most common screening method in women’s health, there are ongoing challenges, including exposure to ionizing radiation, missed malignancies (false negatives), false positives, patient distress and allergic reactions to the contrast agents (Løberg et al., 2015). Moreover, while radiography and ultrasound show anatomical formations, thermal imaging is based on thermodynamics and thus can encompass cellular metabolic discrepancies that produce specific heat responses (Etehadtavakol & Ng, 2017). Thermography has been proposed as a tool for breast cancer monitoring; however, advanced thermographic imaging generally depends on industrial software, which restricts its processing for medical diagnostic applications (Fernández-Cuevas et al., 2015). The development of Dynamic Tele Thermometry Techniques (DAT) enabled time scanning of dynamic areas, maximising the potential of thermal imaging. (Anbar et al., 2000) and reducing false positive and false negative results in relation to the static thermography (Gonzalez-Hernandez et al., 2019; González et al., 2019). Based on DAT, video thermometry (VTM) has been developed as an alternative diagnostic tool that involves acquiring multiple consecutive images at intervals of 15-150ms to monitor dynamic temperature variations (Salhab et al., 2005). High resolution VTM devices take advance of new infrared sensors, which sensitivity has been improved from 0.5°C in the 1960s to less than 0.02°C nowadays, which is effective in capturing subtle temperature variations in breast tissues (Kandlikar et al., 2017). The VTM analysis used in this study process images captured in real-time, optimized for detecting minimal temperature fluctuations in a centesimal range (MART software™ US11931130B2 patent), which potential has been tested for clinical applications (e.g., Cadena et al, 2024). At the laboratory scale, calorimetric systems are employed to measure heat capacity, specific heat and enthalpy changes associated with the cellular metabolism. Calorimetric kinetic studies on ion pump coupling activities have shown that a significant amount of energy released during ATP hydrolysis can be lost as heat instead of being fully converted into electrochemical gradients (de Meis et al., 2005). Heat generated by tumour cells was also associated with the sarcoplasmic reticulum Ca^2+^-ATPase activity in HeLa cells (Suzuki et al., 2007). Thus, it is tempting to speculate that other ion pumps differentially expressed in tumour cells can also significantly contribute to heat release and may serve as new biomarkers for cancer detection.

In preclinical and clinical conditions, it is difficult to discriminate the rate of metabolically heat released by different physiological and pathophysiological phenomena, such as from infections, inflammation, ischemia, allergic reactions, neovascularization, and malignant tumours (Cadena et al, 2024). However, it has been shown that even rapidly growing and poorly vascularized small tumours can raise the regional temperature by 1 to 3°C (Xie et al., 2004), a range that can readily be detected by thermometry (Kakileti et al., 2017) with higher specificity by analysing different tumour depths (Sadeghi et al., 2019). Nevertheless, the lack of an effective cancer metabolic biomarker has limited progress in thermometry diagnostic imaging.

A differential overexpression of V-ATPases at the tumour cell membranes has been described in a range of cancers (Martinez-Zaguilan et al., 1993; Stransky et al., 2016; Couto-Vieira et al., 2020). Highly invasive breast cancer cells show higher levels of plasma membrane V-ATPase compared to less invasive ones, playing an important role in acidifying the extracellular tumour microenvironment (Hinton et al., 2009; Pérez-Chen et al., 2024). Inhibiting this proton pump strictly reduces the migration and invasive capacity of metastatic cancer cells and concanamycin A is used as a standard V-type proton pump inhibitor (e.g., Cotter et al., 2015; Costa et al., 2018; Martins et al., 2019).

Metabolic reprogramming related to tumour cells hyperproliferative behaviour involves the Warburg effect, a key metabolic hallmark of cancer, consisting in a switching from oxidative phosphorylation to glycolysis for ATP production that releases an excess of H^+^ ions inside the cytosol (Cairns et al., 2011). V-ATPases pumping H^+^ across the plasma membrane of tumour cells contribute to maintaining the cellular pH homeostasis and trigger other adaptive responses (Chen et al., 2024). For instance, this efflux of H^+^ ions also leads to a higher acidification of the extracellular environment activating cathepsins and metalloproteases that degrade the extracellular matrix, facilitating tumour cell invasion and migration into surrounding tissues and distant sites, inducing angiogenesis and metastasis (Gocheva et al., 2007; McCarty et al., 2010).

We used an advanced video thermometry (VTM) for detecting thermal variations associated with tumour heterogeneity in abnormal mammary tissues, aiming to characterise thermal signatures that may support preclinical analyses and more precise definition of invaded areas. Moreover, a proof of concept was demonstrated measuring concanamycin-sensitive V-ATPase activity on tumour cell membranes as a biomarker for cancer tumorigenic/ metastatic activities as previously postulated by some of us (e.g., Costa et al., 2018; Martins et al., 2019; Couto-Vieira et al., 2020). This work introduces a new approach combining thermometry using a VTM technique with Scanning Ion-selective Electrode Techniques (SIET) to monitor proton fluxes, alongside spectrophotometric detection of ATP hydrolysis providing a roadmap for the development of more accurate and accessible diagnostics for veterinary and preclinical human health studies.

## MATERIAL AND METHODS

### Patient Selection and Surgical Procedures

For preclinical protocols, the consensus regarding the diagnosis, prognosis, and treatment of canine and feline mammary tumours is followed to standardize the trials (Cassali et al., 2020). Patient selection was based on a comprehensive clinical assessment, including medical history, physical examination, laboratory tests, and imaging studies. Additional criteria encompassed the prescription of supportive care and surgical indication. Postoperative management included continuous monitoring, antibiotic therapy, and pain control.

Six female dogs with mammary tumours underwent total unilateral mastectomy, derived from screening spontaneous tumours at the UENF Veterinary Hospital where the choice was made for the presence of both well differentiated and undifferentiated tumour masses. Incisional biopsies were sent for histopathological diagnosis in the preoperative period. This study was approved by the animal ethics committee ID 432171-CEUA/UENF.

As the exclusion criteria, castration female dogs, female dogs without evident masses of adenomas, carcinomas or mixed variants. As the inclusion criteria, female dog in reproductive phase (standardization of the group to avoid post-castration hormonal variables), without weight or age restrictions, with one or more evident masses of adenomas, carcinomas, or their mixed variants were included in this study.

Surgeries were combined with real-time, non-contact VTM examination, with image captures performed in a temperature-controlled surgical room maintained at 22±2°C (Collins & Ring, 1972; Zhang et al., 2009). In this study, female dogs affected with well delimited differentiated tumours and disperse undifferentiated tumours were analysed. ‘Differentiated tumours’ are understood as organized cells masses composed of solid masses with clear boundaries (Nakagaki et al., 2022). ‘Undifferentiated tumours’ are understood as neoplasms with low structural organization, composed of solid masses with amorphous boundaries (Benesch et al., 2022; Lim et al., 2023). All analysed tumours were submitted to histopathological diagnosis. Three distinct samples were obtained from each patient using VTM: (A) ‘Tumour mass’, (B) ‘Tumour Periphery’, representing transition areas, and (C) ‘Tumour-free area’. Biopsies of control, tumour-free areas were conducted in regions without signs of atypia or tumour activity, and where normal homogenous thermal signals were detected at a distance around both differentiated and undifferentiated tumours. Biopsies from each region (A, B, and C) underwent multidisciplinary analysis, encompassing histological, biochemical and biophysical assessments, integrating thermography and V-ATPase enzyme and proton transport activities.

### Anaesthetic procedure

The veterinary patients received a pre-anaesthetic protocol based on chlorpromazine, at a dose of 0,5 mg/kg intramuscularly in the semimembranosus muscle. Venous access allowed ringer-lactate infusion through with a controlled dose of 10 mL/kg/h. Anaesthetic induction was performed with the association of propofol at a dose of 3 mg/kg, followed by morphine sulphate at a dose 0,4 mg/kg, ketamine hydrochloride at an analgesic dose of 1 mg/kg and lidocaine hydrochloride at a dose of 30 mg/kg.

The animals were then intubated and maintained in a surgical anaesthetic plane using the technique of intravenous partial anaesthesia with continuous intravenous infusion of morphine sulphate at a dose 1,66 μg/kg/min, ketamine hydrochloride at an analgesic dose of 15 μg/kg/min and lidocaine hydrochloride at a dose of 30 μg/kg/min was administered. Isoflurane was administered in an anaesthetic circuit without gas rebreathing for patients weighing less than 7 kg and a semi-closed anaesthetic circuit (with partial gas rebreathing) for patients weighing more than 7 kg. The animals remained on spontaneous ventilation throughout the operation.

During the procedure, monitoring was performed using a multiparametric monitor (Digi care Life Window LW9x Vet®, United States), the heart rate (HR) respiratory rate (RR), electrocardiogram, plethysmography, end-tidal carbon dioxide concentration (EtCO2), oxyhemoglobin saturation (SPO₂) and the blood pressure was measurement using a non-invasive method.

Postoperative analgesia was administered using a multimodal approach, consisting of tramadol hydrochloride at a dose 4 mg/kg, carprofen at a dose 2.2 mg/kg, and dipyrone at a dose 25 mg/kg, all administered orally every 12 hours for five consecutive days.

### VTM analysis

In the preoperative inspection by VTM, the software shows several mini screens with different pseudo-colour gradients (Figure 1), generated by automatic image analysis, where the operator chooses the screen that presents the best image and transfers it to the processing area following the MART’s protocol: 1) assessment of symmetry: considering the small difference in skin temperatures between the right and left side of the homeotherms as around 0.24°C (Uematsu, 1986); 2) vascular inspection: visualization of the main tumour irrigation and the direction of blood flow in the mammary chains; 3) analysis of thermal discrepancy in nearby pixels in areas suspected of abnormalities: Thermal asymmetries result from functional changes and a difference above 0.3°C is suggestive of abnormality (Goodman et al., 1986), while above 1°C is indicative of dysfunction (DiBenedetto et al., 2002); 4) Texture and lesion extent identification: This step involves analyzing grey-scale images formed by subtle temperature variations to reveal textures and boundaries, not measuring temperatures. 5) Detection of similar radiometric thermal patterns: when the operator marks a breast cancer known area and activate the ‘*Similarity tool*’, automatically identifies areas with the same thermal signature, thus enabling the identification of suspicious images to be sample collection. Then, the operator suggests in real time which presumptive samples should be collected by the surgeon during the surgical procedure.

**Figure 1.**
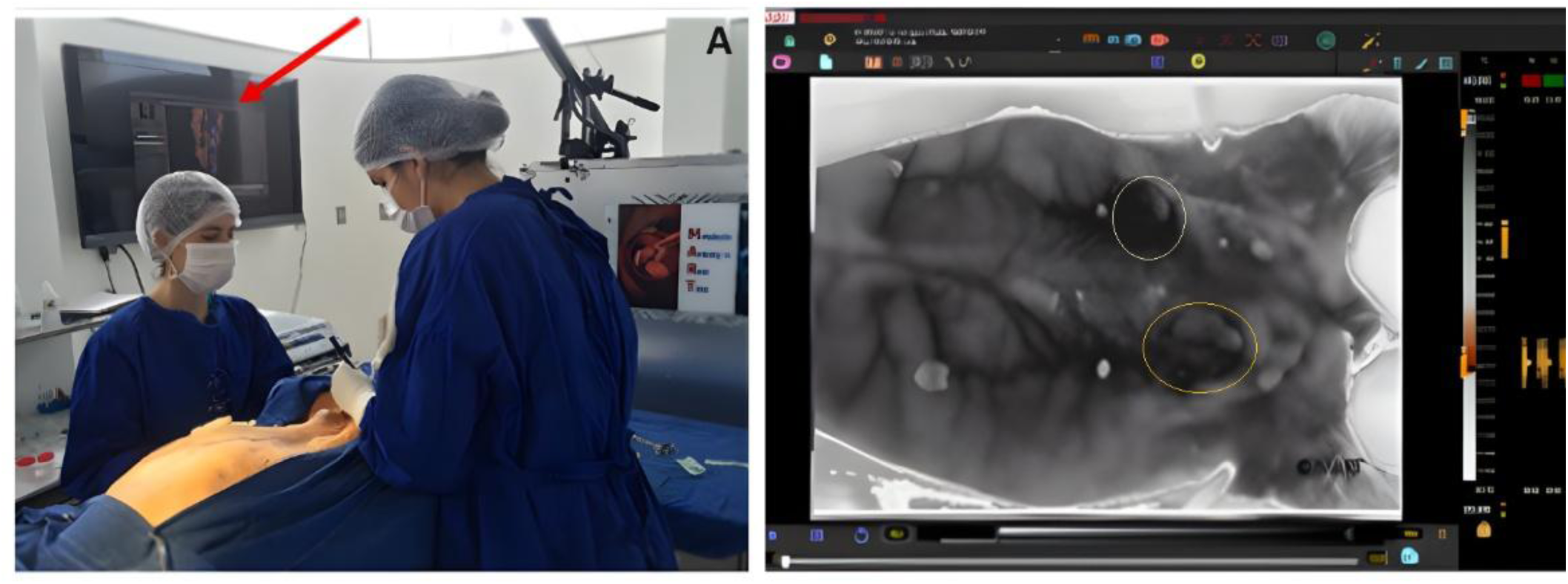
VTM-assisted mastectomy procedure: (a) Surgical room featuring a MART station configured for VTM analysis; (b) Real-time processing results displayed on the surgeon’s monitor, presenting data from a female canine patient diagnosed with mammary cancer.

Grey-scale images are generated by centesimal temperature gradients, which create distinct textures and boundaries. Image clarity is enhanced through the elimination of background noise and exclusion of non-essential regions using the MART ‘Merlin tool.’ This approach allows the selection of pseudo color schemes tailored to various organs and tissues from a palette displayed on MART’s mini screens (Figure 4), enabling the segmentation of only the region of interest as defined by the Merlin tool’s parameters.

### Cell Fractionation Procedure

Biopsied samples were surgically collected guided by the VTM thermal patterns and submitted to differential centrifugation essentially as described by Sennoune and colleagues (2004) with minor modifications (Martins et al., 2019). After surgical removal, the samples were homogenized in a medium containing: 20 mM MOPS-KOH, (pH 7), 200 mM Sucrose, 1 mM PMSF, 1 mM benzamidine, 0.003% BSA, 2mM DTT and 10% Glycerol. Subsequently, the samples were centrifuged following a washing sequence with the washing buffer in a refrigerated centrifuge at 500 rpm for 20 min and 4°C, the process was repeated twice. The precipitate was then homogenized in Poter and the homogenate subjected to a new washing centrifugation at 500 rpm for 20 min. The supernatant was collected and centrifuged in an ultracentrifuge at 100,000 x g for 60 min at 4°C to isolate the microsomal fraction. The pellet was resuspended in a storage buffer containing 250 mM sucrose, 1 mM EGTA and 50 mM Tris-HCl at pH 7.2. The total protein content of the microsomal fraction was determined using the Bradford method (Bradford, 1976), with bovine serum albumin (BSA) employed to the standard curve. Samples were rapidly frozen in liquid nitrogen and stored at -70°C in an ultra-freezer until further analysis.

### Concanamycin-sensitive ATP Hydrolysis Activity

The V-ATPase hydrolytic activity was measured in microsomal membranes derived from cell fractionation of the biopsied samples, in a reaction medium containing: 50 mM Hepes-tris buffer pH 7.0, 5 mM MgSO_4_, 0.2 mM NaMoO_4_, 0.1 M KCl, 0.1mM vanadate and 1 mM ATP. The reaction was started with the addition of each microsomal fractions (0.050 mg/mL of protein) and stopped after 15, 30, 45 and 60 min of incubation, by adding 5% TCA, kept on ice. The phosphate released during ATP hydrolysis was measured using a modified Fiske and Subbarow (1925) method with ammonium molybdate and ascorbic acid (100:1) and read the results at 750 nm on an Elisa reader. The concanamycin-sensitive V-ATPase specific activity was calculated by the difference of ATP hydrolysis with and without 5 nM concanamycin A1 (Martins et al., 2019), which is a highly specific inhibitor of V-type H^+^-ATPases (Huss et al., 2002). Concanamycin A1, a very specific V-ATPase inhibitor, was purchased from Sigma (St. Louis, MO, USA).

### Proton Flux Measurements

Samples were obtained from regions predetermined during the VTM-assisted surgeries. The efflux of H^+^ ions was measured close to the cells using vibrating proton-selective microprobes coupled to a Scanning Ion-selective Electrode Technique (SIET) platform. The signals were obtained from changes in pH that generated electric currents measured by the proton-selective microelectrode.

The SIET data collection captured by 3D microelectrodes that can vibrates on x, y and z axis provide the necessary information to calculate the ion flux of a given point on the surface of a biological sample by means of Fick’s law (J = D (dc/dx)), where D is a tabulated diffusion coefficient for each ion (according to Handbook of Chemistry and Physics, Chemical Rubber Co.). The spatial difference, represented by dx, is derived from calculating the distance between two points defined by the vibration cycle of the microelectrode. Ion-specific currents are measured both near to and at a distance (10–50 micrometres) from the sample surface, and the difference in these currents forms the basis for flux calculation. The concentration difference, represented by dc, is a vector that varies during the assay. At each point the concentration can be calculated from the mV value recorded at the point and the equation and constants determined for the H^+^ specific ionophore during calibration (Ramos et al., 2008).

### Statistical Analysis

Statistical analysis was conducted to compare temperatures and enzyme activity across different sampled regions collected by VTM: (A) Tumour mass, (B) Tumour Periphery and (C) Tumour-free area. A two-way ANOVA test was applied, followed by the Tukey test (p<0.05) for multiple comparisons. Graph Pad Prism version 8.02 was used for these analyses. Principal component analysis (PCA) was performed using Python 3.11 with the scikit-learn library (version 1.3.0).

## RESULTS

Presumptive samples were collected from tumour tissues and surrounding areas for analysis of six female dogs with breast cancer assessed using VTM prior to surgical intervention (Figure 1). Biopsies were guided by VTM thermal signatures characterising three distinct areas: A- Tumour mass, B- Tumour

Periphery and C- Tumour-free area (Figure 2 A and B); following the VTM MART’s protocol: 1) assessment of symmetry; 2) vascular inspection; 3) analysis of thermal discrepancy; 4) identification of textures and lesion extension; and 5) detection of similar radiometric thermal patterns (Figure 3 and 4).

**Figure 2.**
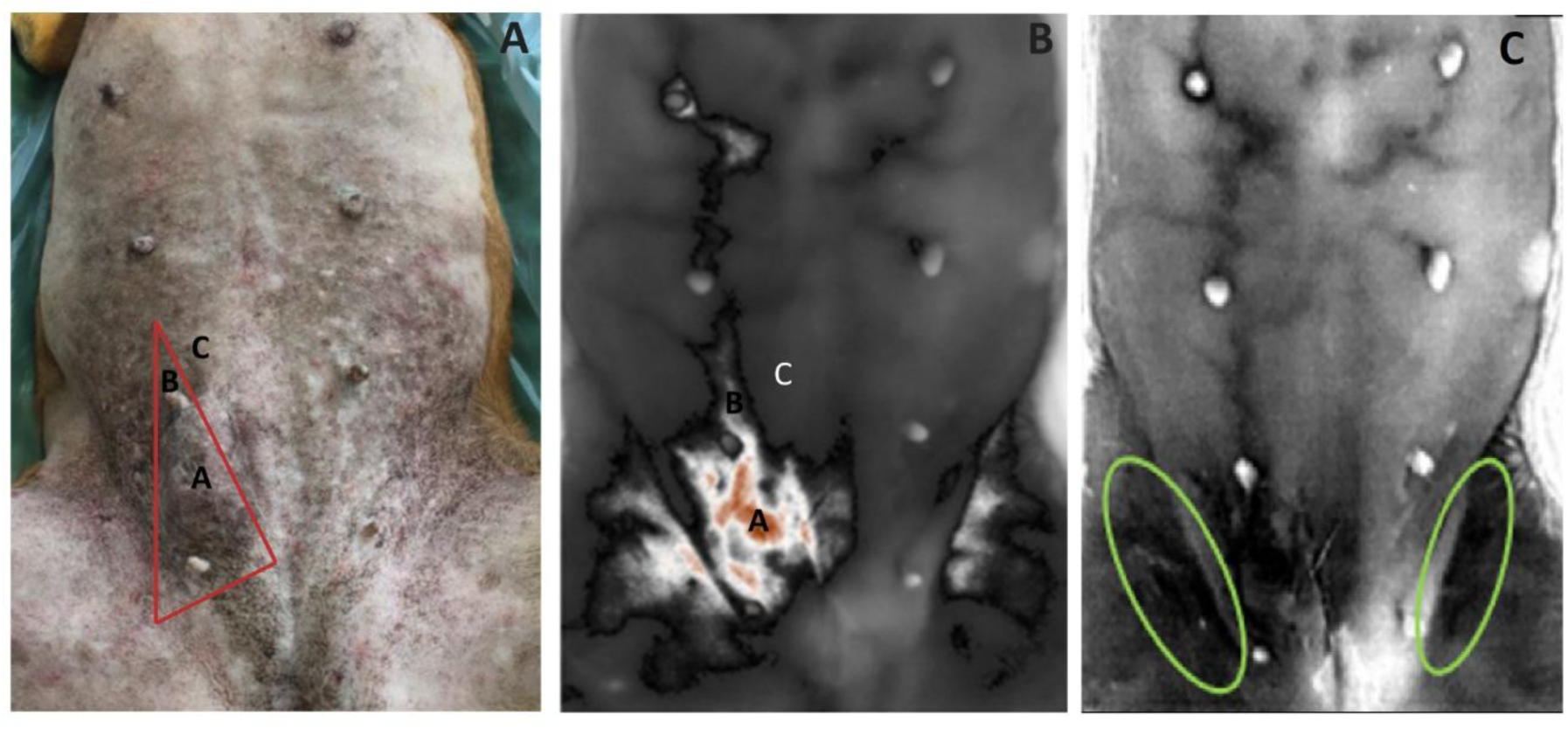
VTM analysis guiding biopsies: (A) Tumour mass with boundaries marked for surgical planning to identify the tumour bed and at-risk tissues; (B) VTM analysis defining areas for sampling biopsies based on thermal patterns in undifferentiated tumours distinguishing, A- Tumour mass, B- Peripheric compromised area, and C- Tumour-free area. (C) The green circles indicate a possible compromised area (on the left) and a typical artifactual signal (on the right) due to depth differences in the inguinal zone.

**Figure 3.**
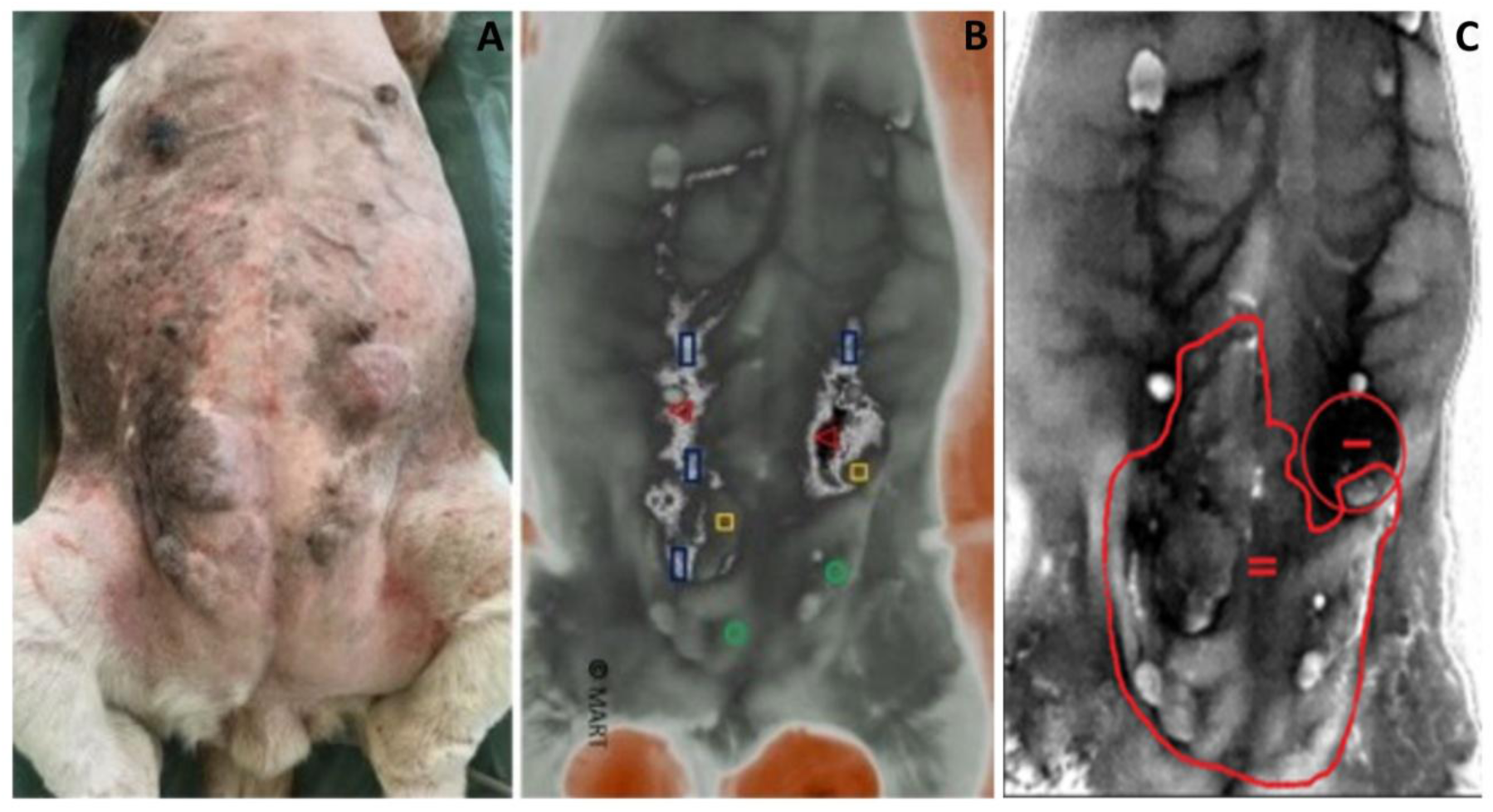
VTM thermal texture and similarity tools: (a) Visible tumoral masses; (b) VTM analysis showing the most thermically active tumours and regions with matching thermal signatures (Red triangles- 38.5oC, Yellow squares- 37.1oC, Blue rectangles- 38oC, green circles- 36.4oC; MART Similarity Tool); (c) VTM vascular inspection indicating well differentiated (I) and undifferentiated (II) tumours in one patient (MART vascular inspection mode - texture search).

**Figure 4.**
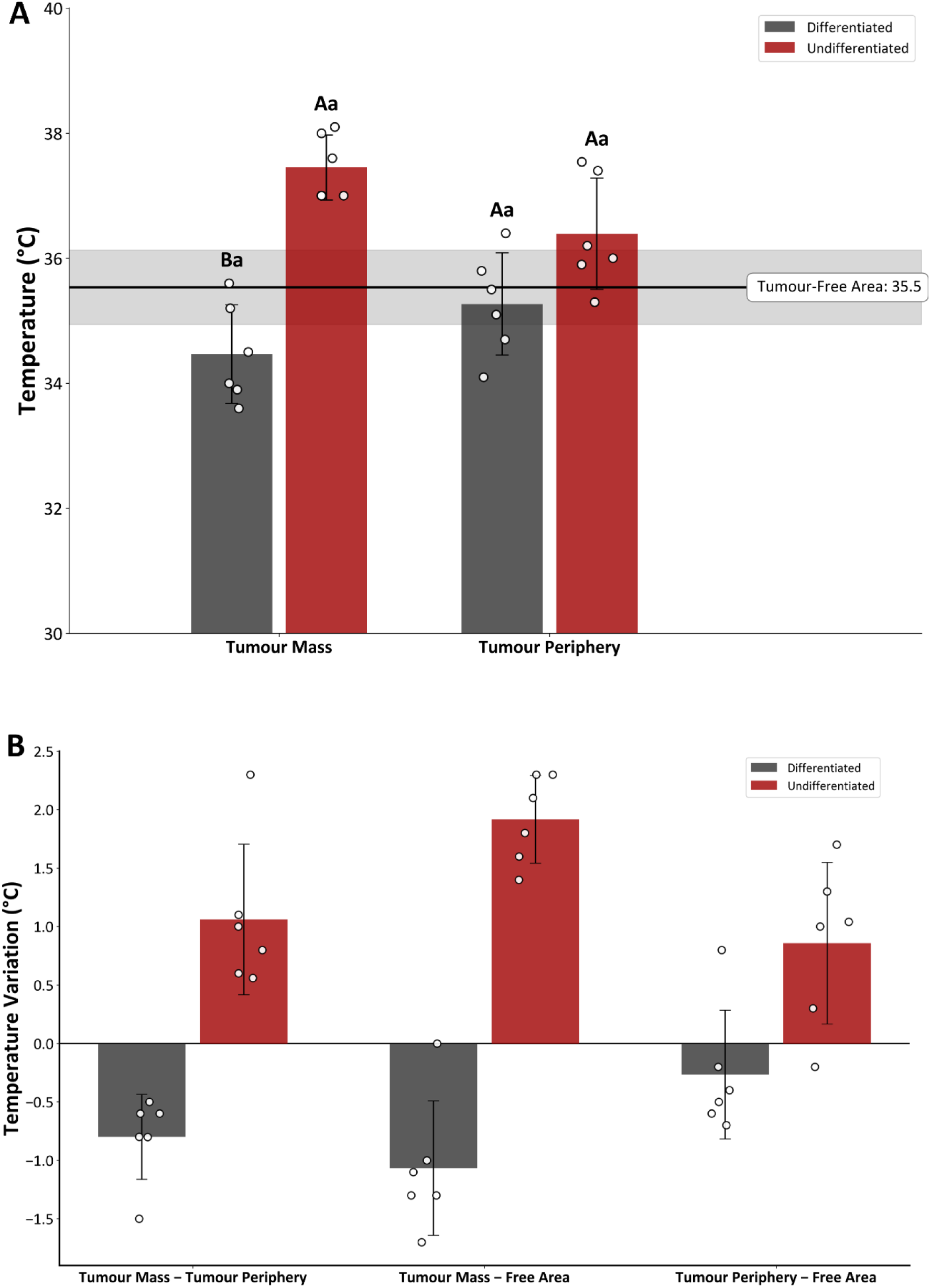
Temperature measurement on three regions of interest: A- Tumour Mass, B- Tumour Periphery, and C- Tumour-free area. (A) Mean temperature (°C) of differentiated (grey) and undifferentiated (red) Tumours Mass, Tumour Periphery and Tumour-Free Area. Different uppercase letters indicate a significant statistical difference between tumour types (differentiated vs. undifferentiated) within the same area. Different lowercase letters indicate a significant difference for the same tumour type between the different areas. (B) Temperature variation (ΔT) between areas A and B (A-B) and between area A and area C (A-C), and between area B and C (B-C) for both tumour types. The bars represent the mean, and the individual points indicate the values for each sample. A two-way ANOVA was performed, followed by the Tukey test (p < 0.05).

Subsequent analyses included histopathological analysis and measurement of concanamycin-sensitive ATP hydrolysis (Figure 5) and H^+^ pumping activities (Figures 6 and 7) used as indicators of tumoral/ metastatic metabolism, followed by evaluation of the integration of all results by exploratory data analysis using PCA (Figure 8).

**Figure 5.**
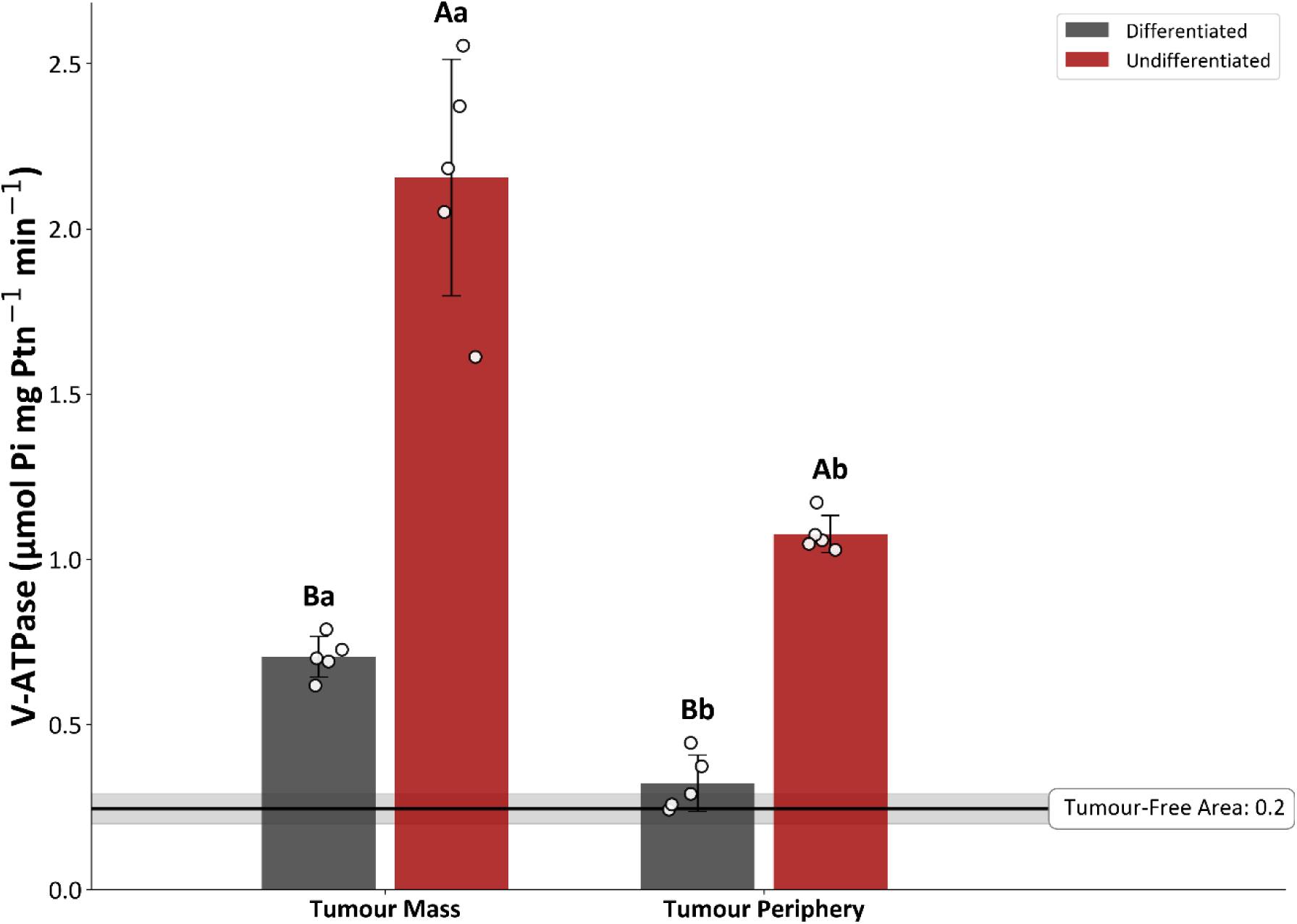
Concanamycin-sensitive V-ATPase enzyme activity: The concanamycin-sensitive ATP hydrolysis of differentiated (grey bars) and undifferentiated (red bars) tumours measured in microsomal membranes derived from: Tumour Mass, Tumour Periphery, and Tumour-free area (shaded line representing a mean +SD activity of health tissues). Different uppercase letters indicate a significant statistical difference between tumour types (differentiated and undifferentiated) within the same area. Different lowercase letters indicate a significant difference for the same tumour type between different areas. A two-way ANOVA was performed, followed by the Tukey test (p < 0.05).

**Figure 6.**
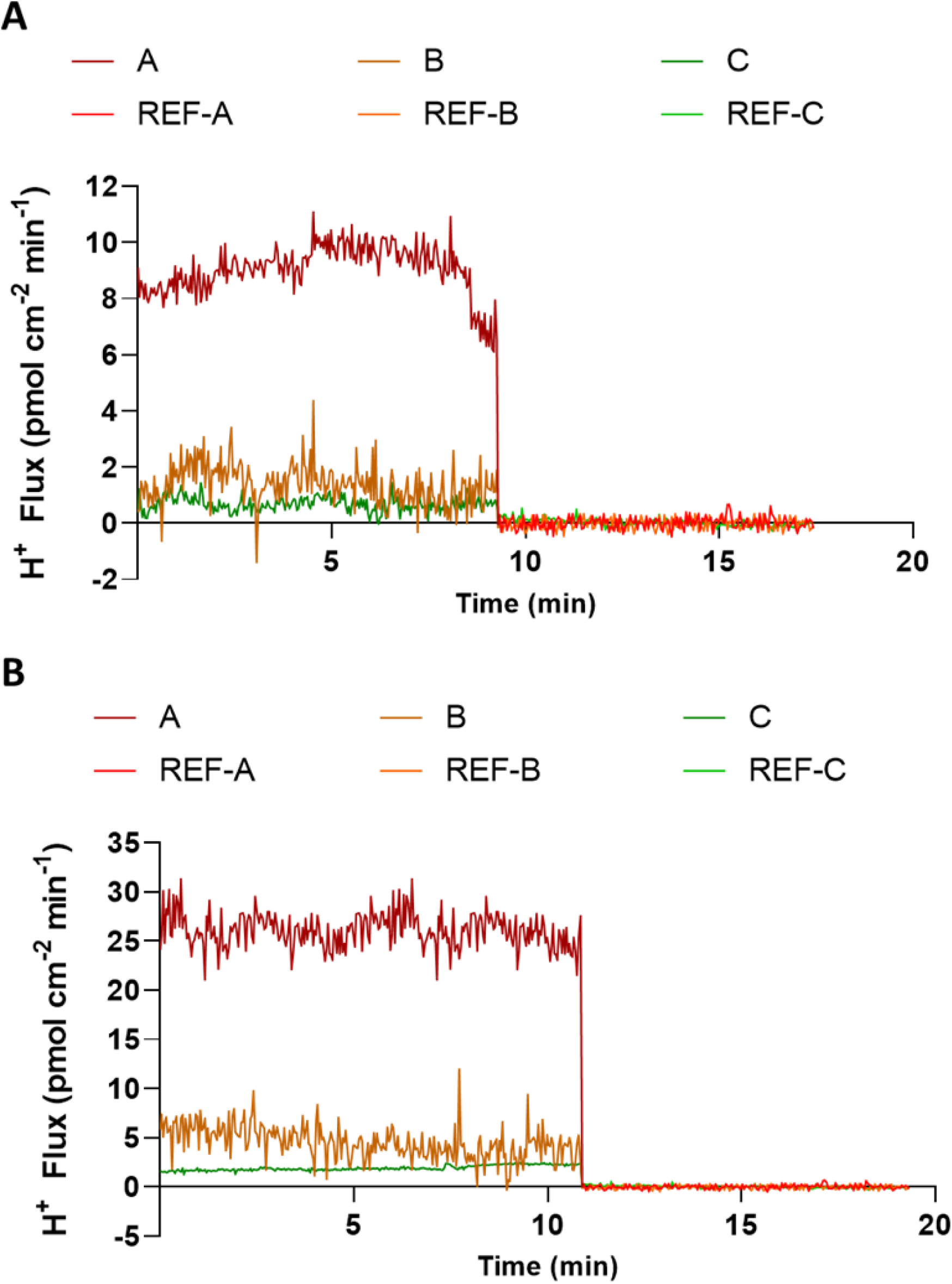
SIET Analysis of proton fluxes: Real-time H^+^ ions flux profiles are shown for (A) differentiated and (B) undifferentiated tumours. Continuous lines represent the dynamic extracellular H^+^ fluxes measured over time. Distinct colours denote the specific sampled regions: Tumour Mass (A, red lines), Tumour Periphery (B, orange lines), and Tumour-free area (C, green lines). Positive values indicate proton efflux, and baseline levels were established by a background reference (Ref.).

**Figure 7.**
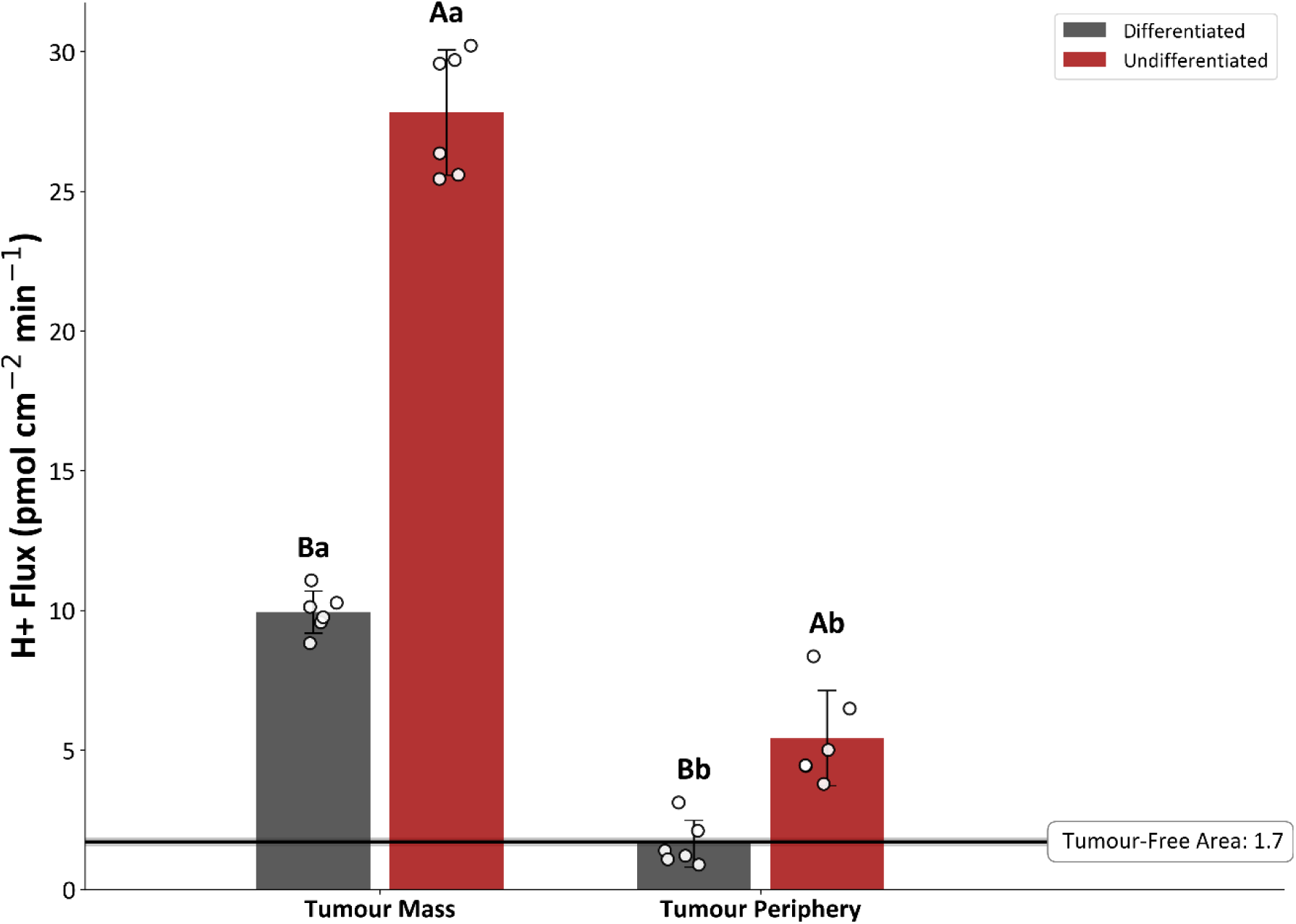
Concanamycin-sensitive proton pumping activity: Bar graphs show the mean H^+^ ions flux for differentiated (grey bars) and undifferentiated (red bars) tumours in Tumour Mass and Tumour Periphery regions. The data is overlaid with individual sample observations (open circles) and error bars. The shaded line indicates the mean baseline H*^+^* flux recorded in the Tumour-free area. Different uppercase letters denote a significant statistical difference between tumour types (differentiated vs. undifferentiated) within the same region. Different lowercase letters indicate a significant difference for the same tumour type between the different regions.

**Figure 8.**
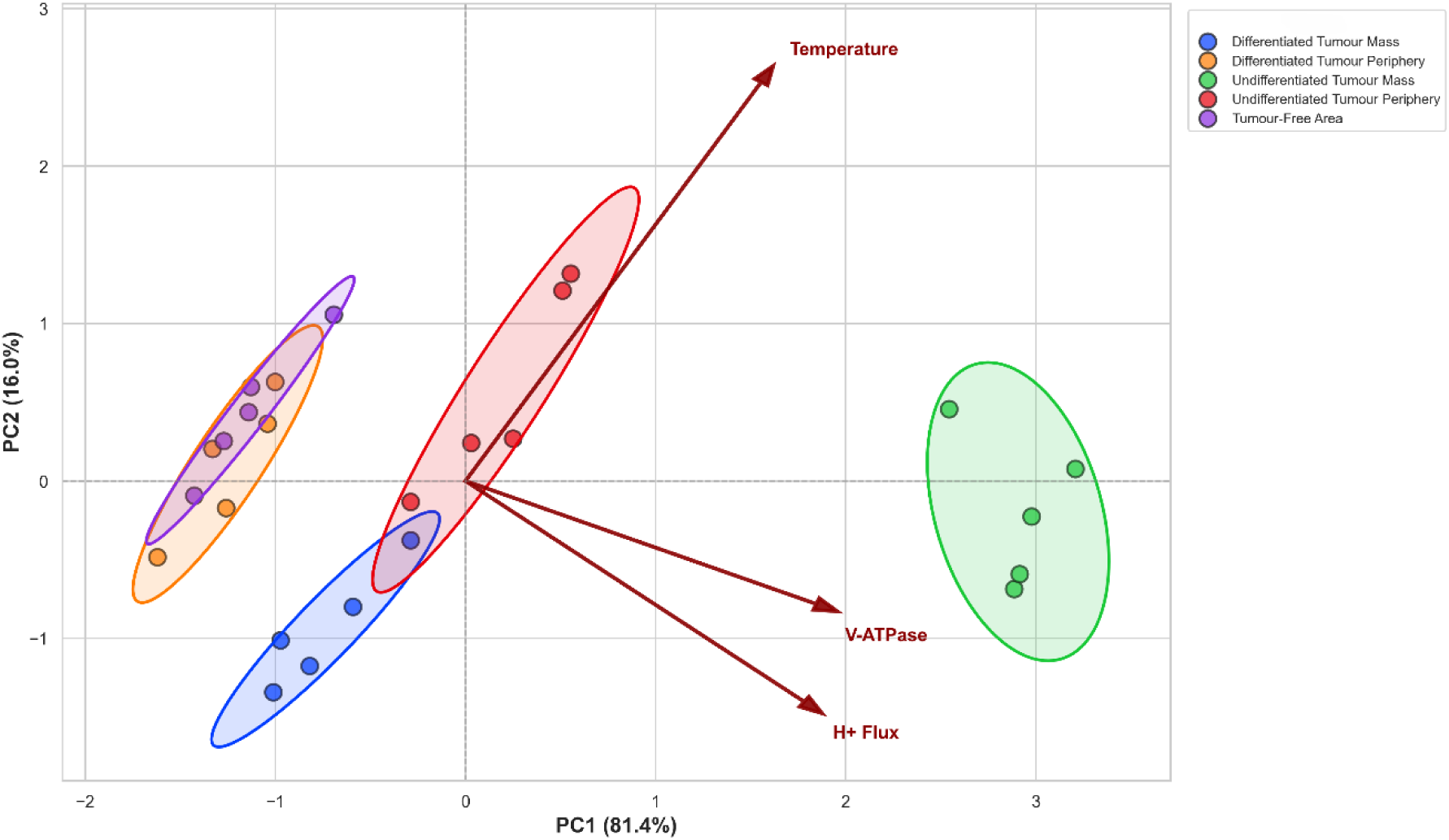
Principal Component Analysis (PCA): The PCA biplot illustrates the relationship between data from differentiated and undifferentiated tumour samples and the tumour-free areas. The first principal component (PC1) explains 91.8% of the total data variance, while the second component (PC2) explains 7.4%. The circles represent the scores for each sample group, and the vectors (arrows) indicate the weight and direction of the contribution (loadings) of each analysed variable. The confidence ellipse highlights the distinct clustering of tumour-free area samples, demonstrating their separation from the other tumour groups based on the measured variables.

### VTM Vascular Inspection by Thermal Imaging Textural Relief

The changes found in the early stages of cancer, such as vasodilation, neo- angiogenesis and high tortuosity of blood vessels are readily recognizable in thermal images (Lahiri et al., 2012). The VTM assessment of symmetry, vascular inspection and identification of thermal textures (Figure 2 and 3) was used to delimitate the lesion extension. The symmetry analysis revealed the tumour masses disturbances on the breast tissue and its vascular supply revealing that the blood supply is large at the base of the differentiated tumour (Figure 3), while the tumour mass has lower mean temperatures (Figure 4). In undifferentiated tumours, vascular network permeates the tumour masses but maintains lower temperatures than those observed at the margin around the area of tumour activity. In the identification of textures and lesion extension, it is possible to see that there are two types of tumour masses: one well-defined tumour bed and another irregular and undifferentiated mass covering a large area (Figure 3C).

### Analysis of Thermal Pattern Discrepancy

Video thermometry regions of interest selected by the MART algorithm revealed significant differences in temperature between tumour bed, margins at risk and tissues free of tumour activity (Figure 2, 3 and 4). This data was used to assist in the mastectomy procedure and to guide sampling biopsies from A- tumour mass core, B- potentially compromised tumour periphery and C- tumour-free control area as defined by VTM analysis protocol. Given the symmetrical skin temperature pattern typically observed between the right and left sides of homeotherms (ΔT < 0.25°C) (Uematsu, 1986) thermal differences greater than 0.3°C can be attributed to functional changes (Goodman et al., 1986; Cadena et al, 2024), whereas differences exceeding 1°C may indicate pathological dysfunction (DiBenedetto et al., 2002; Shao et al., 2023).

Figure 2B shows a routine vascular inspection, where thermal textures appear in the tumour area, but not in the inguinal region where it is possible to observe a potential signal of invasion in the region of contact with the affected region, The green circles in Figures 2C demarcate the inguinal region to highlight the difference between the signal of an at-risk tissue (left circle) from that representing an artefactual signals producing a mirror-image thermal emission (comparing Figures 2B and 2C).

### Analysis of Thermal Pattern Similarities

These representative MART VTM-assisted surgery exemplifies the resolution obtained in VTM analysis. Figure 3 demonstrates comparative VTM tumoral and vascular scanning, which delineates the tumour’s blood supply and identifies principal pathways exhibiting elevated blood flow toward the axillary and inguinal lymph nodes. The vascular thermography mapping provides real-time imaging versatility, facilitating the detection of artifacts, anomalies, and blood flow in regions of interest (Figures 2B and 3B).

Ratiometric VTM analyses revealed thermal patterns similarities by activating the ‘*Similarity tool*’ of the VTM software operator, which are signalised by geometric figures in areas automatically detected as matching thermal signatures selected for sample collection and further analyses (Figure 3B). Specifically in this representative figure ‘red triangles’ highlight a differentiated tumour on the left abdominal mammary line with a corresponding area with a similar thermal signature on the right, both averaging 38.5°C. Signalised by ‘yellow squares’ are similarity zones located inside an undifferentiated tumour, displaying a mean area temperature of 37.1°C. Marked by ‘blue rectangles’, sites along the vascular pathways of the right and left mammary chains, each with an average area temperature of 38°C. Such a bilateral thermal signature along the vascular pathways may indicate potential routes for metastatic spread. Furthermore, in another amorphous regions of this undifferentiated tumour, ‘green circles’ delineate a thermal signature confined to the inguinal regions (right and left), with an average area temperature of 36.4°C.

### VTM Analysis of Tumour Heterogeneity

Tumours bed and peripheric areas at risk were defined by the VTM analysis in high resolution, assessing the distinct thermometric signatures of different type of tumours and areas expressing putative cancer like metabolism (Figure 3B). The MART VTM algorithm enables a delineation of the tumour boundaries by providing thermal images with capacity to reveal tissue surface textures (Figure 3C). Undifferentiated tumours showed significantly higher average temperatures than differentiated tumours (Figure 4A). Although differentiated tumour masses tended to have lower mean temperatures than healthy tissues (by 0.3 to 1°C), the difference was not statistically significant, which may reflect limitations of using mean values to identify underlying tumoral dysfunction (Figure 4B). This may also be related to the harsh metabolic environments and nutritional stresses faced by cancer cells in poor vascularised regions of the tumour (Benzarti et al., 2024).

### V-ATPase Activity - Concanamycin-sensitive ATP Hydrolysis

The V-ATPase hydrolytic activity was estimated by measuring the concanamycin-sensitive ATP hydrolysis in microsomal membranes isolated from biopsied samples from: A- tumour mass, B- peripheric tumour margins and C- tumour-free areas (Figure 5). Differentiated tumours showed significantly lower hydrolytic activities than undifferentiated tumours, in both sampled areas A and B. On the other hand, ATP hydrolysis from thermogenic samples from peripheric area (B) of differentiated tumours showed no significant difference between from control samples (C). Regarding undifferentiated tumours, it was possible to observe a significant difference in V-type (Concanamycin-sensitive) ATPase activity among the three samples, with sample A (tumour masses) showing the greatest activity, followed by sample B (suggesting compromised tumour margins), while sample C (control tumour-free areas) showed the lowest activity (Figure 5).

### V-ATPase Activity - Tumour Cell Proton Flux

Cellular H^+^ effluxes were measured in differentiated and undifferentiated tumour tissues directly from biopsied samples of the tumour mass (A); peripheric potentially compromised tumour margins (B) and control tumour-free areas (C) (Figure 6 and 7). Differentiated tumour masses presented higher H^+^ effluxes than those found in peripheric samples, while tumour-free samples show only a residual H^+^ efflux (Figure 6). Undifferentiated tumours presented a greater H^+^ effluxes than those found in their peripheric samples, with values much higher than that from differentiated tumours. In addition, samples from compromised tumour margins of undifferentiated tumours presented H^+^ effluxes significantly higher than that from healthy tissues (Figure 6).

Proton fluxes from undifferentiated tumours are noticeably higher than that from differentiated tumours, particularly in samples (Figure 7). Conversely, no statistically significant difference was found between the minor H^+^ flux of certain peripheral regions of tumours and those in tumour-free areas, suggesting margins without detectable malignant activity (Figure 7).

### Principal Component Analysis (PCA)

Principal component analysis (PCA) demonstrated that among all the samples assessed, V-ATPase activity and H^+^ flux were predominantly associated with data obtained from undifferentiated tumour masses. In contrast, data derived from the peripheral regions of undifferentiated tumours and differentiated tumour masses were distributed along a central oblique axis in the opposite side of the graph, suggesting the proton pump activities exerted comparatively less influence on the characterization of those biopsied samples (Figure 8). Notably, the temperature vector is located on the same side but in a different quadrant from the V-ATPase biochemical markers, suggesting an indirect correlation between genuine energy dissipative and metabolic energy transduction markers. In line with this notion, data collected from the periphery of differentiated tumors are positioned oppositely in relation to the three eigenvectors, indicating a negative correlation with those bioenergetic markers. This pattern is consistent with cancer-free margins and is further supported by their clustering alongside control samples from tumor-free areas (Figure 8).

## DISCUSSION

Accurately identifying tumour compromised margins is challenging and requires significant expertise, as visual assessment remains the main method in mastectomy and other surgeries used to treat many types of cancer. Although an array of scientific equipment and software can now analyse organs and tissues across a wide electromagnetic spectrum with precise results (Hill et al., 2001; Snyder et al., 2000), there is still an urgent need for effective adaptive tools to improve and validate the specificity of these technologies in preclinical studies before they can effectively be used in clinical practice.

The present work employed a VTM technology derived from intelligent imaging and database-assisted detection methods originally developed for military applications between 1960 and 1990 (Irvine, 2002; Diakides & Bronzino, 2007). It is important to clarify the distinction of this thermometric method from the conventional ‘thermography’. The last analyses static images, whereas the early, generally referred to as ’thermometry’, enables real-time analysis of dynamic environments. Despite, thermometry received FDA approval in 1982 as an adjunctive tool for breast cancer diagnosis, computational constraints until 1989 led to the discontinuation of most clinical applications, although some research activity persisted (Irvine, 2002). Ongoing advancements in high-precision quantum well infrared photodetector cameras and software have led to renewed interest in real-time digital thermometry within scientific and medical fields (Fauci et al., 2001).

Since then, even static thermograms have been proposed as reliable indicators of increased risk of breast cancer by revealing surface temperature changes induced by malignant tumours (Parisk et al., 2003; Arul et al., 2021; Khan & Arora, 2021). Despite further studies investigating infrared diagnosis have approved the sensitivity of the new thermometric devices, their specificity has demonstrated considerable variability (Vreugdenburg et al., 2013; Kolarić et al., 2013; Gogoi et al., 2019). Nevertheless, it has been shown that integrating dynamic states from distinct patterns of changes in breast temperature associated with cold stress it is possible to obtain a thermometric evaluation with greater specificity and precision (Jiang et al., 2010).

The VTM technique employed in this study offered real-time data, delineating tissue textures of tumours and surrounding margins, effectively distinguishing infrared-detected patterns within both differentiated solid tumours that are contained in one area and undifferentiated multiform tumour masses (Figures 2 and 3). On the one hand, the lack of significant differences between data from the edges of differentiated tumours and tumour-free tissues (Figure 4) is consistent with the general notion that surgeries work best for solid tumours that are contained in one area. On the other hand, this may underscore the ongoing limitation of VTM analysis in consistently differentiate areas at risk of cancer from other benign physiological thermal changes (Cadena et al, 2024). Thermal artifacts that can be interpreted as a false positive have been reported during real-time dynamic analysis, as movement can disturb the time-temperature series of each pixel (Riyahi-Alam et al., 2015), an issue that have been addressed by integrating the thermal image with numerical simulations (Singh & Singh, 2020).

The present VTM analysis demonstrated that it is possible to delimit the tumour margin through thermal textures (Figures 2 and 3) and successfully identified thermal patterns correlated with distinctive metabolic heterogeneities (Figures 5, 6, 7 and 8). This agrees with previous evidence that thermometric analyses can achieve much greater precision than the most usual thermography and other diagnostic techniques (Jawzal & Ekici, 2018; Fauci et al., 2001).

Tumour regions with poor glucose supply, due to uneven vasculature, create conditions where only metabolically adaptable cells persist and spread. Cancer cells are defined by this metabolic flexibility, which enables them to adapt and proliferate (Benzarti et al., 2024). It has been postulated that cancer cells can produce more heat than normal cells due to their elevated metabolic activity, which can be at least partially associated with different ion pumps activities and coupling (Suzuki et al., 2007). However, physiological heat releases have also been associated with specific functional couplings of the energy metabolism in highly catabolic normal cells and tissues, primarily linked to activations of F- and P-type ion pumps (de Meis et al., 2005; Hayek et al., 2019). On the other hand, V-type H^+^ pumps have been reported to be differentially overexpressed in the tumour cell membranes, leading to acidification of the extracellular environment and facilitating tumour proliferation and invasion (Hayek et al., 2019). This study presents the first evidence that V-ATPase activity constitutes a distinct component of tumour-derived dissipative energy, which correlates with thermal fingerprints identified through VTM analysis. This relationship is further illustrated by the PCA analysis (Figure 8), where vectors for thermometry and V-ATPase activity data are oriented on the same side of the PCA plot, yet extend toward opposite quadrants, as expected for the coupling between energy dissipation and energy driven reactions in a same transduction system (de Meis et al., 2005).

VTM-selected tumoral samples exhibited clearly distinct proton pump activities, as detected by both the non-invasive Scanning Ion-selective Electrode Technique (SIET) for monitoring proton fluxes and ATP hydrolysis measurements from cell fractionation (Figures 5, 6 and 7). Both activities related to V-type proton pumps as demonstrated by the sensitivity to the specific inhibitor concanamycin A, which exert their specific inhibitory potential at nanomolar concentrations (Dröse et al., 1993; Huss et al., 2002). These parameters showed variability in relation to different tumorigenic activities as expected considering the variation of V-ATPase subunits differentially expressed in different breast tumours (Cotter et al., 2015; Hinton et al., 2009; Sennoune et al., 2004) and composing multi-cancer molecular signatures (Couto-Vieira et al, 2020).

Tumour cells have an accelerated metabolism and are adapted to meet the demands of uncontrolled growth and proliferation (Hayek et al., 2019). The high rate of cell division and the increase in the synthesis of biomolecules higher energy production is required, which implies in increased metabolic activity and released heat as well (Zhu & Thompson, 2019). Here, undifferentiated tumours exhibited the highest levels of both thermal and proton pump activities (Figures 5, 6 and 7). Conversely, although differentiated tumours showed V-ATPase activity higher than that of the control group, their thermometric averages were below normal body temperature reflecting the complexity of tumour heterogeneity and the related diversity of metabolic reprogramming (Couto-Vieira et al, 2020).

V-ATPases are proton pumps present in endomembrane of all eukaryotic cells but that can be found in plasma membranes of some specialised cells playing critical functions in the regulation of ion homeostasis and cell metabolism (Stransky et al., 2016). Overexpression of V-ATPases on the plasma membrane are implicated in many types of tumours creating an acidic extracellular environment that promotes cell proliferation, migration and invasion of adjacent and distant sites, while also maintaining a slightly alkaline intracellular pH that supports tumour growth (Martinez-Zaguilan et al., 1993; Webb et al., 2011). Higher levels of plasma membrane V-ATPase have also been correlated with increased risk of metastasis and poorer prognosis (Capecci & Forgac, 2013). Targeting these pumps with inhibitors is being explored as a therapeutic strategy to impairing tumour aggressiveness by disturbing angiogenesis, extracellular matrix degradation, and chemotherapy resistance (Neri & Supuran, 2011; Martins et al., 2019; Santos-Pereira et al., 2021).

This work demonstrates novel use for concanamycin and any other highly specific inhibitor of V-type proton pumps, in revealing the activity of cancer cell V-ATPases, which serve as a biomarker for tumorigenesis and metastatic processes. Indeed, ion pumps play a crucial role in thermogenesis by actively transporting ions integrating the energy transduction system of cell membranes, from which uncoupling cycles can result in heat generation. Despite extensive research on mitochondrial bioenergetic dysfunctions in cancer, there is a gap in regarding the role of V-ATPase activity in the energy metabolism and thermogenesis associated with tumorigenesis (Luo et al., 2020). As far as we know, the present results provide first evidence demonstrating that changes integrating differential V-ATPase activation and thermogenesis can serve as reliable markers for anomalous metabolic activity related with tumorigenesis and metastasis. This is in line with the notion that the V-ATPase by regulating the acidic microenvironment triggers adaptive responses to the tumour cells increased metabolic activity (McGuire et al., 2016). This metabolic reprogramming subsequently affects tumour behaviour and aggressiveness by enhancing angiogenesis, facilitating extracellular matrix degradation, and enabling tumour cells to adapt, survive, and proliferate in acidic environments that would be inhospitable to normal cells (Pérez- Sayáns et al., 2009).

Taken together these results highlight a close relationship between the tissues H^+^ ion flux and cancer-related reprogramming metabolism that can be used as a marker for more active tumorigenesis and metastatic potential, refining the specificity of VTM and other diagnostic techniques. Furthermore, gaining insight into the distinct metabolic properties of cancer cells is essential for advancing effective therapeutic strategies. This research offers a robust methodology for pinpointing metabolic limitations that may be addressed by innovative treatments. This kind of integrated metabolic analysis has the potential to facilitate the development of more effective and personalized cancer therapies with reduced adverse effects (Okahashi et al., 2025).

## CONCLUSION

The results demonstrate that VTM was able to safely and promptly detect regions potentially affected by tumour dissemination in real time. The data also reveal limitations of this VTM technique in clearly delineating compromised margins with unequivocal metastatic activity. However, this advanced VTM proved effective in distinguishing tumour tissues with varying metabolic activities, synergically, both validated by and validating of, the use of concanamycin- sensitive V-ATPase activity as a biochemical marker for cancer metabolic reprogramming. This body of evidence offers seminal structural correlations with complimentary patterns between thermal and metabolic cues that can establish a robust database for refining VTM algorithms, enabling more accurate and safer identification of tumorigenesis and metastatic activity. The integrated use of thermometry and biochemical-electrophysiological classic techniques open new avenues for cancer research, preclinical trials and preoperative planning in mammary/ breast surgical procedures, potentially with broader applicability to other therapeutic interventions in different types of cancer (Shao et al., 2023).

## AUTHORSHIP CONTRIBUTION STATEMENT

P.G.A.C. (Conceptualization, data curation and analysis, manuscript writing and VTM-assisted surgery methodology); S.B.S (Conceptualization, data curation and analysis, manuscript writing and electrophysiological methodology); B.X.M. (Conceptualization, data curation and analysis, manuscript writing and biochemical methodology); S.O.P.S, S.M.R.C. and T.M.B. (Veterinary medicine and surgery); R.F.A., R.M.S. (VTM interface with biochemical methodology); H.J.L. (Histopathological methodology); S.P.F.C. (Data organization for curation), F.A., G.D.A.B. and L.M.M. (Anaesthesiology methodology), A.L.A.O. (Conceptualization, data curation, manuscript revision and mentoring of the veterinary medicine and VTM-assisted surgery team); A.R.F. (Conceptualization, data curation and analysis, manuscript writing, final revision, and mentoring of the biochemical and electrophysiological team).

## ACKNOWLEDGEMENTS

The authors thank CAPES (001), CNPq (PQ 313221/2025-1), and FAPERJ (CNE E-26/204.302) for financial support to the research groups of A.R.F. and A.L.A.O.; José Edgard O. Alves (PGSA/UEA-UENF) for technical assistance with surgeries; Anna L.O. Façanha (LFBM/UEA-UENF) for critical review and suggestions; and the Instituto Galzu, Rio de Janeiro, Brazil, for providing the MART station.

